# fCCAC: functional canonical correlation analysis to evaluate covariance between nucleic acid sequencing datasets

**DOI:** 10.1101/060780

**Authors:** Pedro Madrigal

## Abstract

**Summary:** Computational evaluation of variability across DNA or RNA sequencing datasets is a crucial step in genomic science, as it allows both to evaluate the reproducibility across biological or technical replicates, and to compare different datasets to identify their potential correlations. Here I present fCCAC, an application of functional canonical correlation analysis to assess covariance of nucleic acid sequencing datasets such as chromatin immunoprecipitation followed by deep sequencing (ChIP-seq). I exemplify how this method can reveal shared covariance between histone modifications and DNA binding proteins, such as the relationship between the H3K4me3 chromatin mark and its epigenetic writers and readers.

**Availability:** R code is publicly available at http://github.com/pmb59/fCCAC/.

## 1 Introduction

Computational assessment of reproducibility across nucleic acid sequencing data is a pivotal component in genomic studies. Moreover, the ever-growing list of available datasets demands robust methods to quickly mine such resources to identify novel potential functional correlations between various genetic and epigenetic regulations. Chromatin immunoprecipitation followed by sequencing, or ChIP-seq, is a widely used method to profile histone modifications (HMs) and transcription factor (TF) binding at genome-wide scale. For each dataset, a set of peaks (regions of statistically significant read counts when compared against an IgG or input DNA controls) can be obtained (Bailey et al., 2013). Reproducibility can then be evaluated by genome-wide Pearson correlation analysis, and peaks in replicates can be compared using Irreproducible Discovery Rate (IDR) analysis and/or overlap analysis (Li et al., 2011; Bailey et al., 2013). However, IDR’s was designed to find a set of reproducible peaks among different replicates of the same type, but cannot be used to compare distinct HMs or TFs datasets. Overlap analysis suffers as well from inherent statistical problems (Bardet et al., 2011). The author has previously developed a methodology that, by using functional principal component analysis, revealed novel correlations between histone modifications that do not colocalize (Madrigal and Krajewski, 2015). Here, I present fCCAC, a functional canonical correlation analysis approach to allow the assesment of: (1) reproducibility of biological or technical replicates analyzing their shared covariance in higher order components; (2) the associations between different datasets. We propose a new statistic to summarize canonical correlations that can be used instead of genome-wide (or peak based) Pearson correlation coefficient, with the advantage of using the profile of the genomic regions to study their covariance at higher orders. We assume that technical and biological replicates will share most of the variability, as will do so *bona-fide* interactions between different co-factors. Overall, fCCAC greatly facilitates the assessment of covariance in genomic applications.

## 2 Implementation

Functional data analysis is a raising field of statistics that allows moving from discrete measurements to functional approximations using an expansion in basis functions (Ramsay and Silverman, 2005). As in Madrigal and Krajewski, 2015 we have used cubic splines to approximate data, which we read from genomic coverages in bigWig format. For N genomic regions (peaks provided in Bed format) we have two sets of curves, (*x_i_*, *y_i_*), *i* = 1,…,N. The curves are then centered, and principal modes of variation ρ_ξ*i*_ and ρ_η*i*_ between *x_i_* and *y_i_* in terms of probe weight functions ξ and η can be estimated (Supp. Material). The N pairs of probe scores represent shared variability if they correlate strongly with one another. Then, squared canonical correlations R_1_^2^, R_2_^2^,…, R_k_^2^, k = 1,…, K, can be calculated as in Ramsay and Silverman, 2005 by constraining successive canonical probe values to be orthogonal. Values of R_k_^2^ close to 1.0 imply high covariation between the two samples. Details of fCCAC operation can be found in Suppl. Information. For K squared canonical correlations, we can compute a weighted squared correlation as 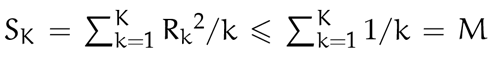, where the weights 1/k are the k-th harmonic number, and decrease with the order of the canonical component. Then, we can report S_k_ as a fraction over the maximum F(%) = 100 × (S_K_/M), where F represents an overall measure of shared covariation. The user interacts with a single function called *fCCAC* (examples can be found in the Supp. Material).

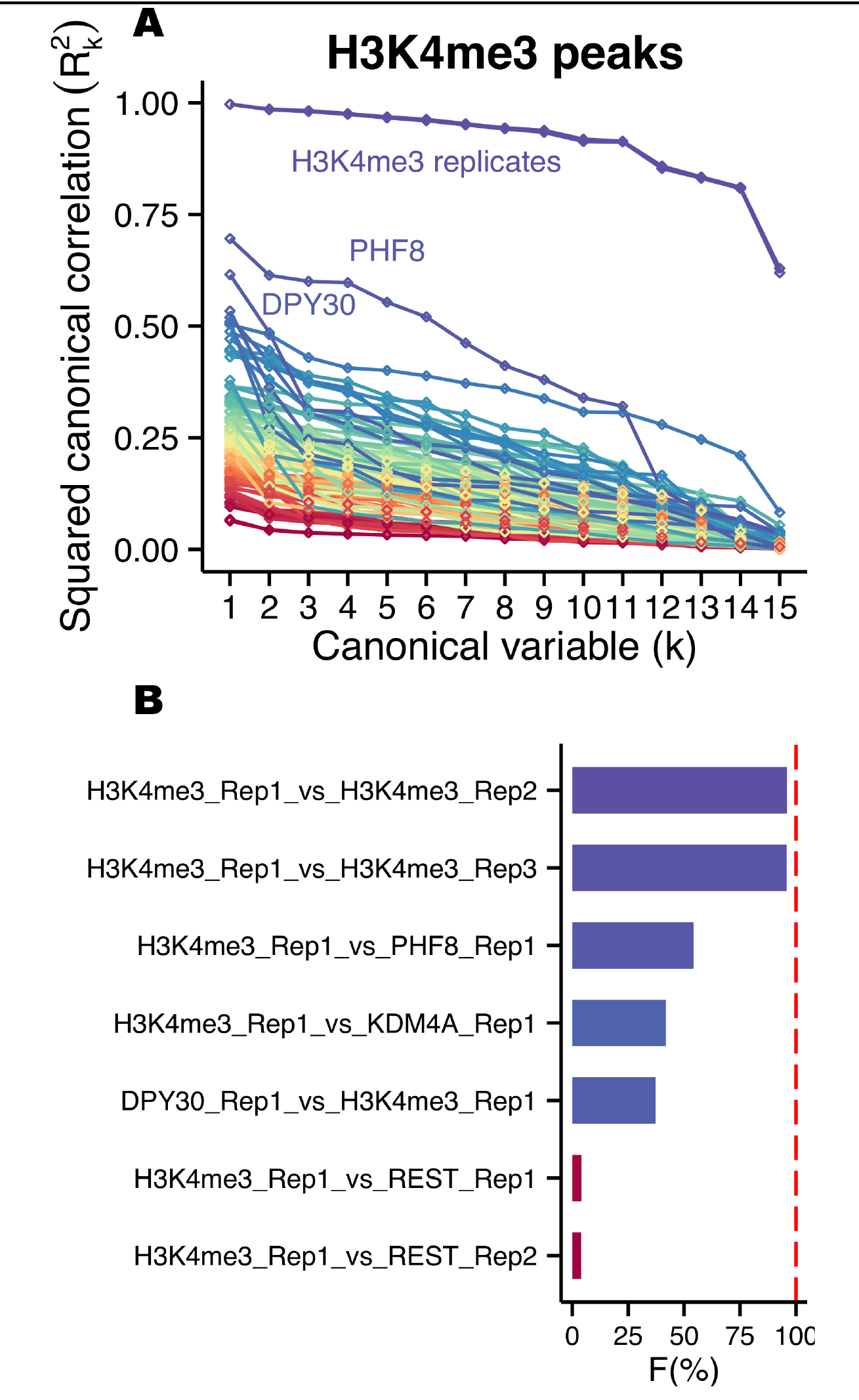
Fig. 1: (A) Squared canonical correlations for H3K4me3 (Rep1) and 59 protein-DNA binding datasets (DPY30 and 58 ENCODE TFs). Spectral colormap based in the value in k = 1. (B) First 5 and last 2 ranked interactions according to their percentage over maximum F. The red dashed line indicates perfect covariance.

## 3 Results

To exemplify the methodology we explored the correlation between the nucleosomal HM H3K4me3 (trimethylation of lysine 4 on the tail of histone 3) and several TFs and chromatin epigenetic remodelers.For this, we focused on human embryonic stem cells (hESCs). We took advantage of recently published H3K4me3 ChIP-seq data (Bertero et al., 2015), which was performed in biological triplicate from the H9 hESC line. First, we defined an aggregated list of peaks at H3K4me3 as our reference set to study replicate reproducibility (23,422 peaks). The results showed high shared covariation (F >95%) for the H3K4me3 ChIP-seq triplicates, as expected (analogous analysis for H3K27me3 confirmed the irreproducibility of one of the replicates; Supp. Material). Then, we analyzed the relationships between H3K4me3 deposition and other genomic datasets for DNA binding proteins. For this, we included ChIP-seq data for DPY30 (Bertero et al., 2015), since this protein is part of the enzymatic complex responsible for the deposition of the H3K4me3 mark, as well 58 other DNA binding proteins included in the ENCODE dataset for the H1 hESC line (97 datasets) (ENCODE Project Consortium et al., 2012). We found high canonical correlations between H3K4me3 and DPY30 (Figure 1A), as expected (Bertero et al., 2015). Only PHF8 (F=54.2%) and KDM4A (JMJD2C) showed higher F value than DPY30 (F=37.2% Figure 1B), in agreement with their known ability to bind to H3K4me3 (Feng et al., 2010; Pedersen et al., 2014). When we monitored all possible combinations of interactions in H3K4me3 regions, TFs BRCA1 and CHD2 showed F=92% in H3K4me3, in agreement with motif analyses suggesting that they might form part of the same complex (Kheradpour and Kellis, 2014) (Supp. Material).

## 4 Conclusion

FCCAC represents a more sophisticated approach when compared to overlap analyses and Pearson correlation of genomic coverage. This method can be used (1) to evaluate reproducibility, and flag datasets showing low canonical correlations; (2) or to investigate covariation betweeen genetic and epigenetic regulations, in order to infer their potential functional correlations. Overall, this method will facilitate the development of new hypothesis regarding how TFs, chromatin remodelling enzymes, histone marks, RNA binding protein, and epitranscriptome changes can cooperatively dictate the specification of cell function and identity.

## Acknowledgements

The author would like to thank Alessandro Bertero for useful comments in the manuscript.

